# Standardized Protocols for Global and Local Thermal Nociception in Black Soldier Fly (*Hermetia illucens;* Diptera: Stratiomyidae) Larvae

**DOI:** 10.64898/2026.05.23.727392

**Authors:** SO Durosaro, M Barrett

## Abstract

Nociception, the capacity to detect tissue-damaging stimuli such as noxious chemicals or high heat, is increasingly studied across insect orders and life stages, informing our understanding of its adaptive value and molecular mechanisms. The black soldier fly (*Hermetia illucens*; Diptera: Stratiomyidae) is widely recognized as the star of the growing insects as food and feed industry. Black soldier fly larvae (BSFL) are reared at high densities that, combined with their extraordinary metabolism, can generate lethal overheating on farms. Data on the thermal nocifensive capabilities of BSFL could inform our understanding of larval behaviors during overheating events (and potential welfare impacts of thermal slaughter), and provide a comparative datapoint to the well-studied vinegar fly (*Drosophila melanogaster;* Diptera: Drosophilidae). Accordingly, we adapted global and local thermal nociception methods from larval vinegar flies for use with BSFL. We find that global assays (akin to boiling) adapt easily to both first and late instar BSFL, that BSFL exhibit slightly different nocifensive behaviors than last instar vinegar fly larvae, and that BSFL have higher thresholds (thrashing begins at 39.70 °C in late instar BSFL versus 26.6 °C in *D. melanogaster*). In contrast, the classical local nociception assay did not adapt easily to BSFL, so we modified the assay to test a time-bounded, continuous exposure to the temperature probe instead of response-latency. This assay generated a unique, gradual pattern of increasing responsiveness to the probe in late instar BSFL (>95% responsiveness above 66 °C) rather than the sharp cutoff in responsiveness at 52 °C demonstrated in *D. melanogaster* during response-latency assays. Altogether, these protocols open the door for standardized research on BSFL thermal nociception for fundamental and applied purposes.

## Introduction

*Hermetia illucens* (Diptera: Stratiomyidae), also known as the black soldier fly (BSF), has exploded in popularity as a research species in the last twenty years (Rumbos and Athanassiou 2021). This increase in study is directly related to its newly discovered agricultural potential: BSF larvae (BSFL) can reclaim many waste streams through bioconversion into useable protein and generally require less water, energy, and land than vertebrate livestock to rear, making them a key part of the future of sustainable and circular agriculture (van Huis 2013; van Huis and Tomberlin 2017; Chia et al. 2019; Purkayastha and Sarkar 2022; van Huis and Gasco 2023; Lalander et al. 2025). Trillions of BSFL are now reared each year around the globe, and by 2033, they are projected to be the most farmed agriculture species for food and feed (McKay and Shah 2025).

Recent reviews of the neurobiological and behavioural literature (Gibbons et al. 2022; Barrett and Fischer 2025) have generated a growing consensus that there is a realistic possibility of sentience (the capacity for subjective experience, with negatively or positively valenced states) in at least some insects (Low 2012; Andrews et al. 2024; Zipple et al. 2024). According to common frameworks used in these reviews, evidence for sentience is strongest in the Diptera, albeit much weaker at the juvenile compared to the adult life stage (partly due to a paucity of studies on juveniles; Gibbons et al. 2022). As a result of the uncertainty about insect sentience, many ethicists, insect producers, and entomologists working with this novel livestock farming industry have called for a precautionary approach (Birch 2017) that favors further research into farmed insect welfare, including for black soldier flies (de Goede et al. 2013; IPIFF 2019; Loon and Bovenkerk 2021; van Huis 2021; Barrett et al. 2023; Klobučar and Fisher 2023; Kortsmit et al. 2023; Andrews et al. 2024). As in other agricultural species for which sentience is still uncertain, though increasingly expected to be plausible (e.g., fish, shrimp, chickens), research into welfare can help mitigate potential harm, even while research into the fundamental capacity for sentience remains ongoing (Barrett and Fischer 2023).

Pain is considered an especially important part of evaluating the welfare of any livestock animal. For instance, the five freedoms framework specifically calls for “freedom from pain” (Brambell 1965), and pain is one of the negative mental experiences that should be avoided within the Five Domains framework (Mellor and Reid 1994). Nociception, defined here as the capacity to sense and avoid tissue-damaging stimuli, can contribute to the development of the feeling of pain in animals that have the capacity for subjective experience (Tracey 2017). Therefore, creating standardized nociception assays for BSFL to understand what stimuli they detect as noxious will be a necessary first step for understanding the welfare of BSFL in relation to the possibility of pain. While standardized protocols for testing nocifensive behaviors exist in *Drosophila melanogaster* (Diptera: Drosophilidae) vinegar fly larvae for a host of noxious stimuli (including heat, cold, chemicals, and mechanical stimuli; Im and Galko 2012), there is significant variation in behavior, morphology, and development between the vinegar fly and black soldier fly. Species-specific protocols are thus warranted to adapt the established vinegar fly nocifensive assays for BSFL.

Accordingly, this study aimed to translate global and local thermal nociception assays currently used for *D. melanogaster* larvae (Chattopadhyay et al. 2012) for use in BSFL. Global assays for vinegar fly larvae involve immersing them in water and heating the water rapidly to assess their responses to noxious temperatures. Local assays involve stimulating a portion of the larval body with a high-temperature probe and assessing their responses. Developing these assays for BSFL has both fundamental and applied value. Fundamentally, this will allow for further investigation into the neurobiology of nociception in BSFL for comparative purposes as, to date, the study of insect nociception has been limited to a handful of model systems, particularly at the juvenile life stage (Gibbons et al. 2022).

Developing these methods for BSFL is also relevant in an applied setting, as it may increase our understanding any negatively-valenced experiences faced by BSFL during two welfare challenges identified in the insects as food and feed industry: 1) heating (e.g., boiling, baking, steaming) as a method of slaughter; and 2) lethal overheating due to high stocking densities (Barrett et al. 2023). In regards to slaughter, understanding the total potential duration of ‘suffering’ experienced by larvae during heating means that it must be clear when environments transition from innocuous to noxious temperatures during the slaughter process (Almeida et al. 2026, Sánchez-Velázquez and Hernández-Álvarez, 2026). Further, identifying a standard suite of nocifensive behaviors performed by larvae would allow for real-time welfare assessment by producers uncertain if their slaughter SOP is potentially causing pain-like states.

For the second, BSFL generate extraordinary metabolic heat during the bioconversion phase; high stocking densities combined with wet, insulating substrates can result in significant heat retention (Barrett et al., 2023; Li et al. 2023). For instance, in-substrate temperatures can be 10 - 20 °C above ambient due to larval metabolic heat production (Li et al. 2023) and stressful temperatures (even when sublethal) can negatively impact production through growth and development impacts (Schow-Madsen et al. 2025). Little is known about how BSFL may sense stressful or lethal temperatures and how their behavior shifts in response to sublethal-but-noxious temperature stress (Barrett et al. 2023). However, improved understanding of larval nocifensive behaviors could allow for production optimization and welfare improvements by giving farmers specific behaviors and noxious temperature thresholds to look for *prior to the onset of lethality* that would predict a lethal larval overheating event is imminent.

We thus used the protocol developed in Chattopadhyay et al. (2012) for testing global thermal nociception in both first (L1) and late (L5/6) instar BSFL. Given significant differences in late instar body size between BSFL and vinegar fly larvae, we describe a modified version of the local nociception protocols designed for use with late instar BSFL only. Using our protocol, we described BSFL behaviors in response to global heat as well as the latency of these behaviors, which differed both from *D. melanogaster* and also by instar in BSFL. Finally, we also described and provided videos of late instar BSFL behaviors in response to local heat, showing how increasing temperature shapes nocifensive rolling responses.

## Methods

### Obtaining Larvae

0-24 hour old larvae (first instar, L1; n = 97) for the global nociception assay were obtained by purchasing eggs from Evo Conversion Systems (College Station, TX). We collected larvae within the first 24 hours of the eggs hatching on day four post-lay. The body mass of first instar larvae was assumed to be that of an egg (25 μg; Dortmans et al. 2017). Late instar larvae for the global thermal nociception assay (L5/6; n = 97) were obtained from Evo Conversion Systems at this life stage. Late instar larvae weighed 0.06 ± 0.005 g (range: 0.05 – 0.07 g) and only lightly colored, non-sclerotized larvae (not prepupae) were used.

Late instar larvae for the local thermal nociception assay (L5/6; n = 864) were obtained from Fluker Farms (Port Allen, LA) at this life stage. They weighed 0.17 ± 0.05 g. Late instar BSFL weights are highly variable (0.05 – 0.40 g) based on nutrition, temperature, and stocking density.

### L1 and L5/6 Global Thermal Nociception Assays

For L1 black soldier flies, we followed the global thermal nociception procedure by Chattopadhyay et al. (2012) for *D. melanogaster*, with minor adaptations. Briefly, their procedure involves putting a water drop on a petri dish, placing the larva in the drop, placing the petri dish onto a heated plate, and then observing five stereotyped behaviors (head thrash, roll, whip, seizure, and paralysis).

For our assay in BSFL, we set a hotplate (Onilab MS-H280-Pro) into the visual field of a Leica L2 microscope, with fiber optic light guides to ensure adequate contrast for viewing the larva at 2.5X magnification. Lights were LEDs to avoid potential ambient heat. We set the hotplate to 60 °C for at least 15 minutes, and confirmed the surface temperature using a thermologger (Omega HH506A) and thermocouple (Physitemp MT29-1HT) before beginning the assays. We placed a 50 μL droplet of water in the middle of a custom PTFE plate (2.5 by 3.1 cm, and 1.75 mm thick) wrapped with Rosco matte black Cinefoil on the dorsal surface and sides only. We ran ten trials recording the temperature of the water droplet after the PTFE plate was placed on the heat plate using the thermologger and thermocouple (until the water reached 60 °C), allowing us to determine the heating rate of the water droplet (Supplementary Figure 1). We also tested a subset of larvae (n = 10) in room temperature water (23.2 °C) that we did not heat for two minutes, to see what behaviors could be attributed to being in water versus the heating of the water.

An Alex 000 paintbrush was used to gently transfer larvae from the rearing container into the water droplet. Within ten seconds of transfer, the PTFE plate containing the larva and water droplet was placed onto the hot plate surface and the timer was started. Assay start time, and the onset of each behavior (move, contract and hold, curl start, curl stop, seizure, and paralysis) were recorded using a voice recorder (Google Pixel 7) to avoid looking away from the larva and later transcribed onto a datasheet.

A generally similar process was used for L5/6 (n = 97, mean ± SD = 0.0626 ± 0.0053 g), with minor adaptations as their greater strength and size allowed them to escape water droplets on the PTFE plate. Therefore, we changed the water droplet volume to 1400 uL, which was placed in a 4 dram glass vial (note that BSFL can grow even larger than those we used, as BSFL are highly variable in body mass at the final instar; if replicating our method with larger larvae, this would require a larger water volume and vial). The hot plate set temperature was changed to 115 °C, and the hotplate itself was covered with a small piece of Cinefoil to increase contrast with the larva for viewing. As in the prior assay, larvae were placed in the room temperature water and then moved to the hot plate and the recording started. We recorded water temperature after vial introduction until the water reached 60 °C (∼2 minute trials) to get a heating curve to estimate water temperature. In this case, larval behaviors were able to be recorded via video (Google Pixel 7) due to their larger size and scored after the assay was complete. We also tested a subset of larvae (n = 10, mean ± SD = 0.0704 ± 0.0052 g) in room temperature water (23.6

°C) that we did not heat for two minutes, to see what behaviors could be attributed to being in water versus the heating of the water.

### Late Instar Larval Localized Heat Assay

We initially attempted to replicate the response-latency assay procedure from Chattopadhyay et al. (2012) for both L1 and L5/6. Briefly, they use a custom-built thermal probe to deliver a localized (0.07 mm^2^ area) heat stimulus to 3rd instar vinegar fly larvae on the dorsal midline of abdominal segment A4 for up to 20 s, or when the larva rolls laterally by at least 360°, whichever comes sooner. Response latency is the recorded output. We were unable to replicate the procedure with first instar BSFL - the probe was too large and would dry them out before they were able to respond. Therefore, we only tested L5/6 black soldier fly. Due to the large size of our L5/6 black soldier flies compared to *D. melanogaster*, we found that L5/6 black soldier flies were strong enough to simply free themselves from the touch of the probe and move away. Therefore, we modified the assay as described in the next paragraph.

Using a custom-built thermal probe with a small metal pointed tip (0.07 mm^2^ area, 47.11° point angle; ability to be set from room temperature to 70 °C, ProDev Engineering, same model as Chattopadhyay et al. 2012), we applied the probe to the dorsal midline of the junction between the second and third segment of the BSFL. The probe was applied perpendicular to the larvae with enough force to lightly indent the larval epithelium and prevent them from moving away from the probe. The probe was held against the larval epithelium for fifteen seconds; it was then removed, and the rolling response of the larvae (which consisted of one or more 360 degree rolls clockwise or counterclockwise, as in *D. melanogaster* larvae) was recorded. We recorded a total of 96 larvae per treatment temperature, with treatments of ‘room temperature’ (the probe was off; n = 95), 27, 46, 50, 54, 58, 62, 66, and 70 °C (n = 863, mean ± SD = 0.17 ± 0.05 g). The start temperature of 46 °C for assessing the nociceptive threshold was chosen based on results in *D. melanogaster* and pilot tests with BSFL.

### Statistical Analysis

Claude Cowork Opus was used to perform an initial check of the dataset for basic data entry errors, but all analyses and coding were performed by humans. All analyses were conducted in GraphPad Prism v. 9.3.1 or R v. 4.4.2. Alpha was set to 0.05 for all analyses and Shapiro-Wilk normality tests were used to assess normal distributions of the data. To determine the line of best fit for heating rate of the water, a nonlinear regression was used in GraphPad; comparison of fits between a quadratic and cubic model revealed that cubic models were the best fit for both L1 and L5/6 water heating rates. As data on number of rolls during the local heat nociception was not normally distributed, a Kruskal-Wallis test was used to compare medians followed by a Dunn’s MCT to separate treatments. A binomial logistic regression with a likelihood ratio test was used to assess the relationship between temperature and rolling probability in the late larval nociception test, with 22 C used for the room temperature datapoints, using R packages *MASS* v.7.3-61 (Ripley and Venables, 2002), *car* v.3.1-3 and *carData* v.3.0-5 (Fox and Weiseberg, 2019). Supplemental videos, raw data and code for this project can be found at: https://osf.io/adhgp

### Ethics Statement

Although there are no legal mandates for the ethical treatment of insects in research at this time, we applied the principles of the 3Rs to our work. Care was taken to reduce the number of animals used for data collection via pilot testing and an *a priori* power analysis. Additionally, larvae may regain motor function following paralysis in a few cases in the global heat nociception assay according to prior reports in *D. melanogaster* larvae; therefore, each larva was euthanized immediately following the assay via crushing with a heavy weight and then frozen for 24 h at -20 °C. During the global heat assay trials, all unused eggs and larvae were euthanised by crushing and then freezing for 24 h at -20 °C. In the local heat nociception assay, initial knockdown trials with an anesthetic in the laboratory failed; therefore, larvae were checked for a return to normal behavior, then were allowed to pupate and then frozen for 24 hours at -20 °C prior to disposal. In the future, the laboratory would update this protocol for ethical reasons to instead humanely euthanize larvae immediately following the assay to avoid the potential of pain; we would use newly-developed anesthesia protocols (Durosaro et al., data in prep), followed by freezing for 24 hours at -20 °C prior to disposal.

## Results

### Global assay heating rates

For the water droplets used with L1, we obtained an average heating rate of 0.44 ± 0.02 °C/s in the first minute of the test, and 0.26 ± 0.01 °C/s over the full test period (temperature = 3.05 e-05 (time^3^) - 0.0085 (time^2^) + 0.84 (time) + 22.73; F = 925.5, df = 1, 606, R^2^ = 0.99, p < 0.0001). Water droplet start temperature ranged from 22.2 to 23.2 °C (average: 22.82 ± 0.34 °C).

For the water in the glass vial used with L5/6, we obtained an average heating rate of 0.34 ± 0.03 °C/s in the first minute of the test, and 0.30 ± .02 °C/s over the full test period (temperature = -6.49 e-06 (time^3^) + 0.0001 (time^2^) + 0.38 (time) + 22.64; F = 13.38, df = 1, 607, R^2^ =0.98, p = 0.0003). Water droplet start temperature ranged from 22.9 to 25.5 °C (average 24.04 ± 0.86 °C).

### Description of behavioral progressions for unheated controls and heated treatment larvae in the global heating assays

L1 do not exhibit a strong water entry response, but may occasionally curl once when first introduced to the water; this always occurs before the droplet is moved to the hot plate, and thus was never captured in our treatment recordings. L1 do not engage in their normal terrestrial movement patterns when in water, even in the absence of heat (control larvae). Instead, larvae release an air bubble from their most posterior segment, which often causes them to get stuck to either the surface of the water bubble or the foil beneath them. When this occurs, they move their whole body, presumably in an attempt to free themselves, in an undulating manner that closely resembles a very slow ‘air dancer’/’inflatable tube man’. Waiting too long to start the assay in an attempt to minimize this irrelevant movement caused the larvae to remain still even after heating began, and thus perform poorly in the assay (e.g., minimal response, even for seizures/paralysis in treatment larvae). Therefore, it is best to start the assay within ten seconds of placing the larva in the droplet, even if they begin the assay ‘stuck’.

When observing control L1 behavior for two minutes after being placed in the unheated water droplet, they moved, were ‘stuck’, or exhibited very slow transitions into the C or U shaped posture - resembling a curl but without the rapidity of contraction and release in a true ‘curl’ as described below. Often, this occurs with the animal first lifting its head and then its tail, instead of rapidly contracting both sections towards one another. Further, the head and tail do not touch and often remain much further apart than a true ‘curl’ as described below.

Treatment L1 subject to heat may complete five behaviors during the assay:

⍰ **L1 Move:** The larva instead begins to move their head in a lateral motion persistently (L1) or undulate their full body like an ‘air dancer’; this movement is usually lacking in urgency (e.g., not a ‘thrash’, and in direct contrast to their behavior when ‘stuck’ for L1), and may include slow, 360 degree rolls. L1 may push their mouthparts against the walls or bottom surface if they can reach it, as though to drag themselves forward. This may also include moving the tip and tail slowly into a C or - more commonly - U shape, but there is no rapid contraction and release like for a curl and the animal may hold this posture for a prolonged period. Often, the tip and tail are held quite far from one another in this posture. This behavior is also exhibited by unheated controls.
⍰ **Contract & Hold:** The larva tightly contracts ventrally, such that their mouthparts are directly touching their most posterior segment. The larva holds that position for at least a second.
⍰ **Curl:** The larva brings their mouthparts and most posterior segments towards one another in a fast contraction and release; if the larva is not stuck with a water bubble, this occurs after the larva has tipped onto its side or dorsal surface. The movement brings both halves of the body into a C or U shape. The mouthparts and most posterior segments do not touch (as in contracting and holding). It must be *fast* (a single, smooth motion happening in 1 s) - slowly orienting in a C or U shape does not count.
⍰ **Seizure:** The larva stretches along the anteroposterior axis, with a maximum of a 30-45 degree bend only to the posterior sections. The larva exhibits a high-frequency, whole body shaking that may be minimal or very violent.
⍰ **Paralysis:** The larva relaxes and stretches along the anteroposterior axis and all movement ceases.

However, several of these behaviors are difficult to reliably observe in L1. The air bubble causes larvae to work to free themselves, and can make it challenging to distinguish the behaviors related to heat v. those related to the posterior end being stuck to a surface in the water and preventing free range of motion (though seizure and paralysis can be reliably assessed even in this case).

Late instar larvae are more active, both in the presence and absence of heat. Many exhibit an entry response, which occurs only in the first ten seconds or less of a larva’s time in the water. As with the L1, it takes this amount of time to move the larva from the set up area to the hot plate; thus, these entry responses were generally not observed/recorded in our treatment animals but we still note them here.

⍰ **Entry response:** In the first ten seconds, larvae respond either to prior handling (which directly preceded entry into the water) or to the water entry itself by rapid side-to-side motion, half-rolls, or full-body rolls as they attempt to right themselves in the water. This subsides within a maximum of 10 seconds following entry into the water, as the animals transition into regular ‘Move’ behaviors, and then through the rest of the behavioral sequence if they are in heated water.

Some larvae do not exhibit the entry response and instead transition directly into ‘move’ behaviors when they enter the water.

Move behaviors for L5/6 larvae can be described as:

⍰ **L5/6 Move:** The larva instead begins to undulate their full body; this movement is usually lacking in urgency, and may include slow, 360 degree rolls. L5/6 may push their mouthparts against the walls or bottom surface as though to drag themselves forward. This may also include moving the tip and tail slowly into an S, C or U shape, but there is no rapid contraction and release like for a curl. This behavior is also exhibited by unheated controls.

As the water heats, L5/6 follow a similar behavioral progression as the L1 but with one additional behavior, Thrash, which occurs in 100% of larvae after movement begins and before the first curl or contraction in the sequence towards death:

⍰ **Thrash:** The larva begins to thrash their body side to side or up and down urgently; this sometimes causes 360 degree rolls that are a consequence of thrashing (but, these rolls should not be taken as an indicator of thrashing alone, see: ‘move’).

Control L5/6 larvae in unheated water will begin the sequence with ‘entry response’ or ‘move’ behaviors and will then continue with ‘move’ behaviors for their entire two minutes in the water. No thrashing, contracting, curling, holding, seizure, or paralysis behaviors were ever observed in control larvae in unheated water; this was directly in contrast to 100% of treatment larvae which exhibited these behaviors from 29 - 114 s after water entry, and suggests that those behaviors are in response to the heat and *not* the prolonged immersion in water. Note that rolling behavior is a consequence of movement in the water and cannot be used to reliably distinguish moving in the water generally from thrashing specifically; therefore, unlike in *D. melanogaster*, rolling on its own is not consistently indicative of thermal nociception in BSFL during a global heat assay.

Overall, this nocifensive sequence is easy to observe in L5/6; supplementary videos of each behavior are provided at https://osf.io/adhgp (Right/Move: Videos 1 and 2; Thrash: Videos 3 and 4; Contract and Hold: Videos 5 and 6; Curl: Videos 7 and 8; Seizure and Paralysis: Videos 9 and 10; Full videos of all behaviors in sequence: Videos 11 - 14). For examples of entry response behaviors without subsequent nocifensive behaviors in control L5/6 larvae in unheated water, see Videos 15-16.

### Population Responses to Global Noxious Heat in First and Late Instar Larvae

100% of treatment L1 moved, seized, and became paralyzed during the progression to lethality (n = 97); 90% of larvae (n = 87) curled at any point during the progression. In 77% of cases (n = 75), behavior proceeded in order as: move/contract (in any order, but without curls), curls, seizure, paralysis. Although curls may be interspersed with movements or contractions, it is nearly always the last stage before seizure (89% of cases). Moving, contractions and holds, and curling can overlap or occur out of sequence in L1 and curls/contractions/holds can be omitted. Only 7.22% of L1 (n = 7) ever contracted and held; this may be artificially low due to the air bubble issue (but in many cases where larvae were unstuck for the entire assay no contracting or holding occurred). The first nocifensive behavior (curl or contract, exhibited by 89% of larvae) was exhibited at a mean water temperature of 42.69 °C in L1; death, measured by the proxy of paralysis, occurred at a mean water temperature of 51.00 °C (Figure 1).

**Figure 1.**
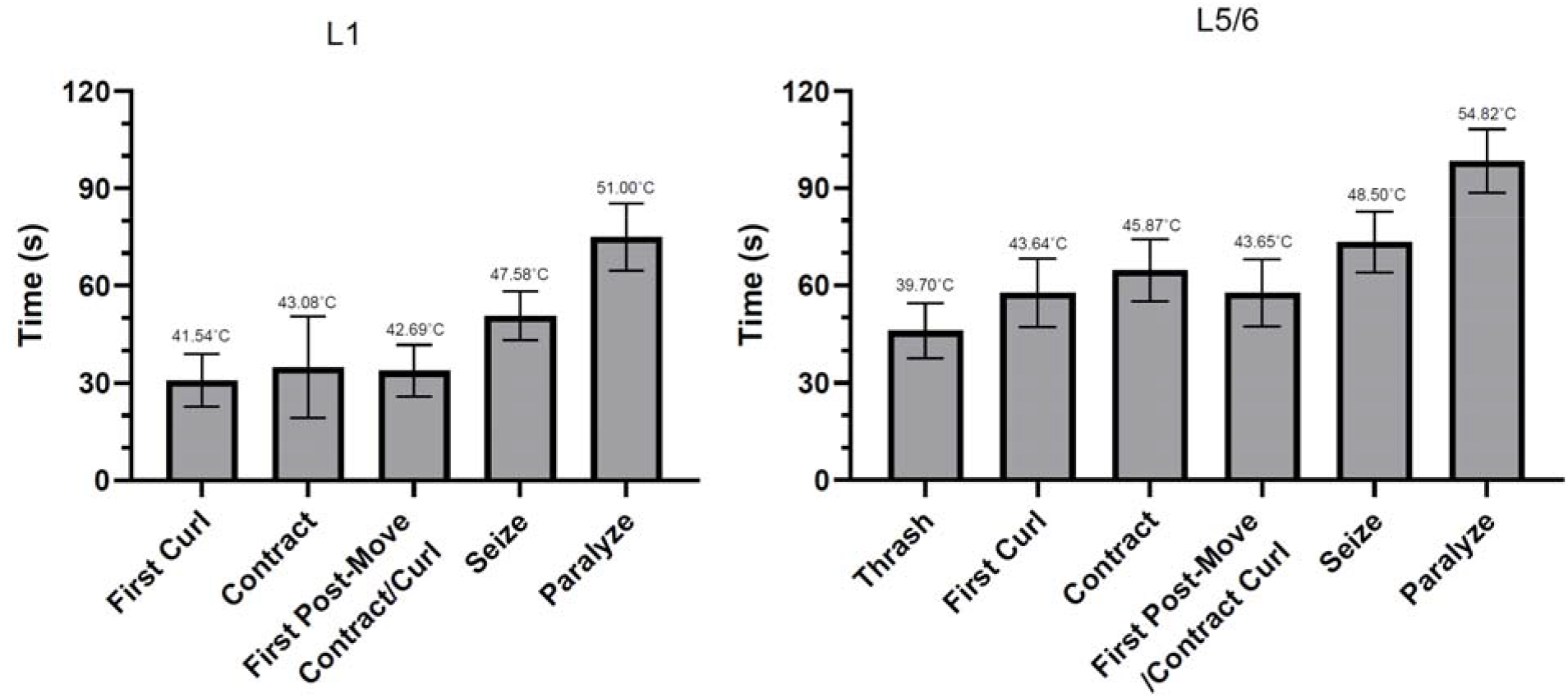
Average latency and estimated water droplet temperatures for each behavioral response at each juvenile life stage for BSFL. The latency to first nocifensive behavior (contract/curl for L1; thrash for L5/6)) and first motor-failure behaviors (seizure and paralysis) were recorded for first instar (L1) and late instar (L5/6) BSF. The estimated water temperature was determined using the formulas obtained in Supplemental Figure 1. Mean ± SD

100% of treatment L5/6 moved, thrashed, seized, and became paralyzed during the progression to lethality (n = 97); 85% of larvae (n = 82) curled at any point during the progression. In 91% of cases (n = 88), behavior proceeded in order as: thrash, curls and/or contractions, seizure, paralysis. Thrashing, contractions and holds, and curling can overlap or occur out of sequence in L5/6, and contractions/holds/curling can be omitted. In contrast to L1, 53% of L5/6 (n = 51) contracted and held. The first nocifensive behavior (thrash, exhibited by 100% of larvae) was exhibited at a mean water temperature of 39.70 °C in L5/6; death, measured by the proxy of paralysis, occurred at a mean water temperature of 54.82 °C (Figure 1). For one individual, the animal seized, briefly curled, and then returned to seizing before paralysis.

L1 exhibited between 0 and 7 total curls while L5/6 exhibited between 0 and 9 total curls (Figure 2).

**Figure 2.**
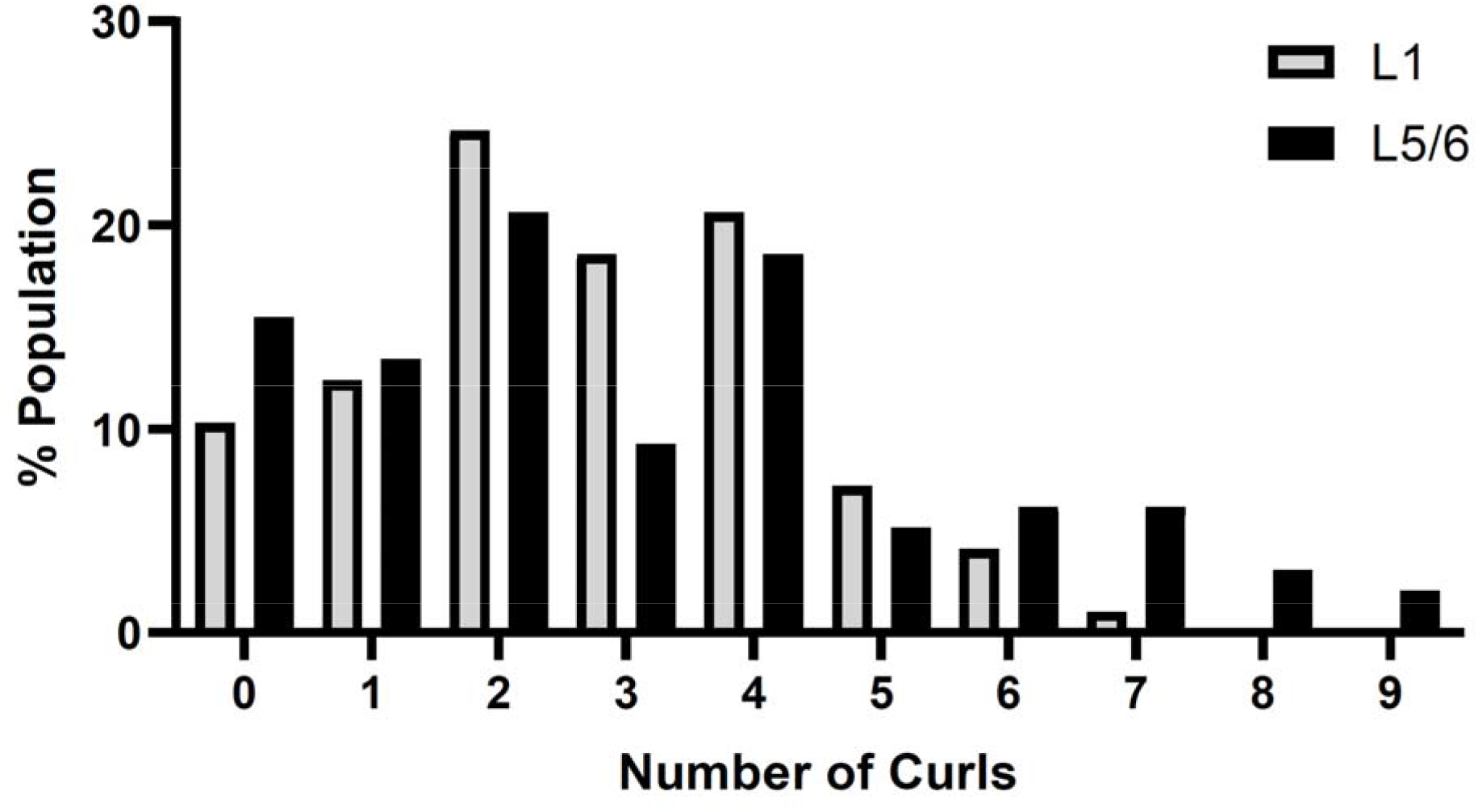
Percent of L1 and L5/6 that curled between 0 and 9 times during the global heat nociception assay. The total number of curls exhibited by L1 was negatively correlated with first curl latency (Figure 3; F = 4.75, df = 1, 85, p = 0.032; y = -1.30 x + 34.74), and positively correlated with seizure onset time (F = 13.62, df = 1, 95, p = 0.0004; y = 1.65 x + 46.20) and the difference between curl and seizure onset timing (F = 19.17, df = 1, 85, p < 0.0001; y = 2.68 x + 12.45). There was a marginally non-significant positive correlation with paralysis onset timing (F = 3.49, df = 1, 95, p = 0.065). L5/6 showed similar trends: a negative correlation between number of curls and first curl latency (F = 6.96, df = 1, 80, p = 0.01; y = -1.39 x + 62.66), a positive correlation with the difference between curl and seizure onset timing (F = 18.83, df = 1, 80, p < 0.0001; y = 1.81 x + 9.94), and marginally non-significant positive correlations with seizure and paralysis onset timing (seizure: F = 3.07, df = 1, 95, p = 0.083; paralysis: F = 3.37, df = 1, 95, p = 0.07). These data suggest that curling is a sustained response to noxious temperatures and not a phasic response occurring only at the onset of noxious temperature perception; because curl count scales with the duration of the window in which curling is possible, it should thus not be used as an index of stimulus intensity or aversiveness.

**Figure 3.**
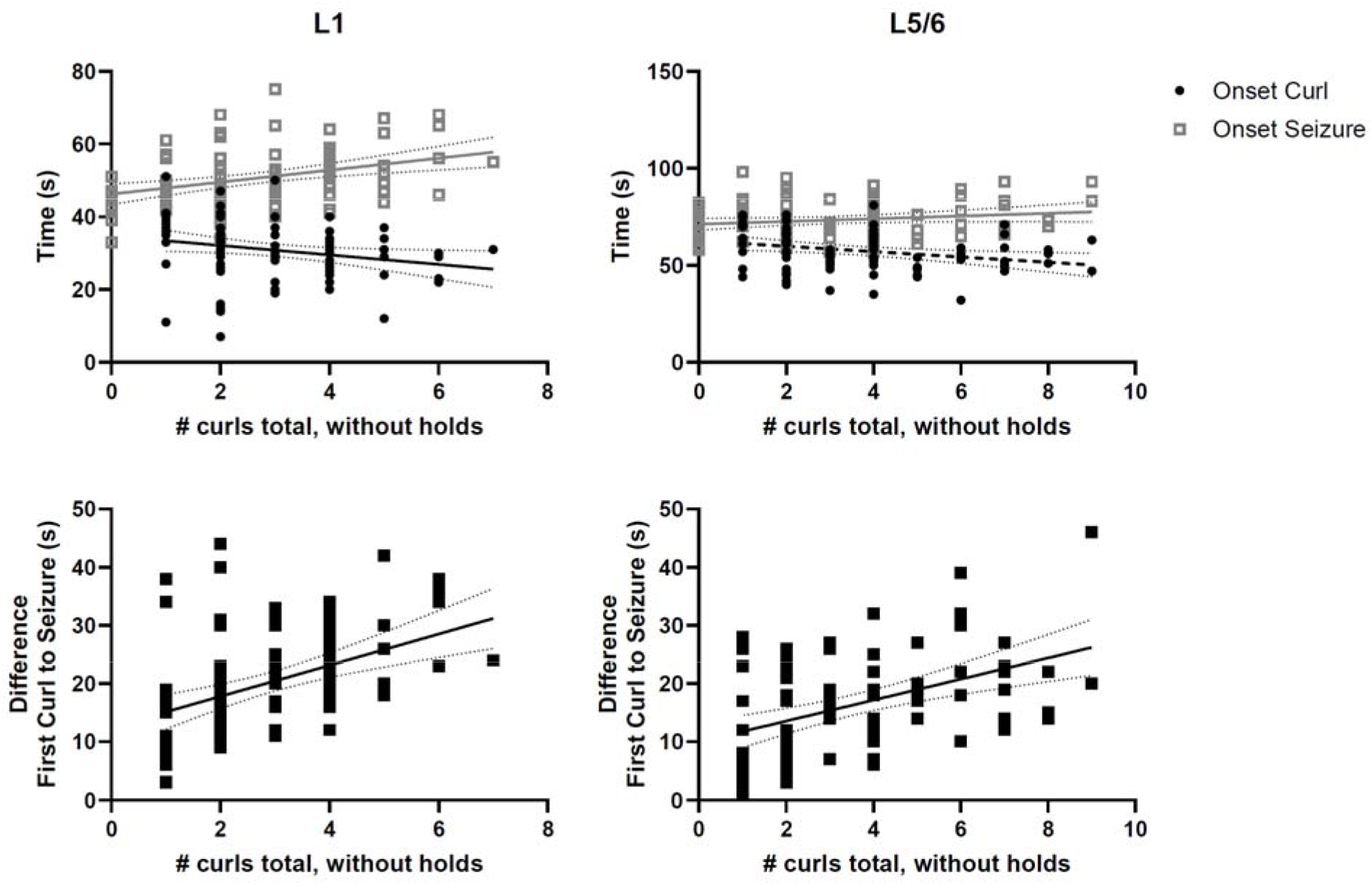
Correlations between the number of curls and the onset timing of curling, seizure, and the difference between curling and seizure onset time.

### Behavioral Progression and Population Responses to Local Noxious Heat in Late Instar Larvae

After the heat probe was removed, L5/6 exhibited the following behaviors:

⍰ **Move -** The larvae began to move normally away from the area where they had been assayed.
⍰ **Turn -** The larvae exhibited a number of 180 degree lateral ‘turns’ or rotations around the anterior-posterior axis; an incomplete ‘turn’ of only 180 degrees was not counted as a roll.
⍰ **Roll -** The larvae exhibited a number of 360 lateral corkscrew rolls (Videos 17-18; https://osf.io/adhgp), similar to nocifensive rolling behaviors observed in other species (e.g., *D. melanogaster* larvae). Rolls were obtained by dividing turns by 2; any incomplete (odd) number was not counted as a roll, meaning animals that turned only once were not counted as rolling.

The number of larvae that rolled was temperature-dependent (Figure 4; Table 1, binomial logistic regression, LRT, X^2^ = 503.45, df = 1, p < 0.0001), with 0-1% rolling at room temperature and 27 °C and 95-96% rolling at least once by 66 - 70 °C. The logistic regression 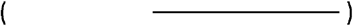 predicted 54.38 ± 0.54 °C as the temperature where 50% of larvae will roll at least once.

**Table 1.**
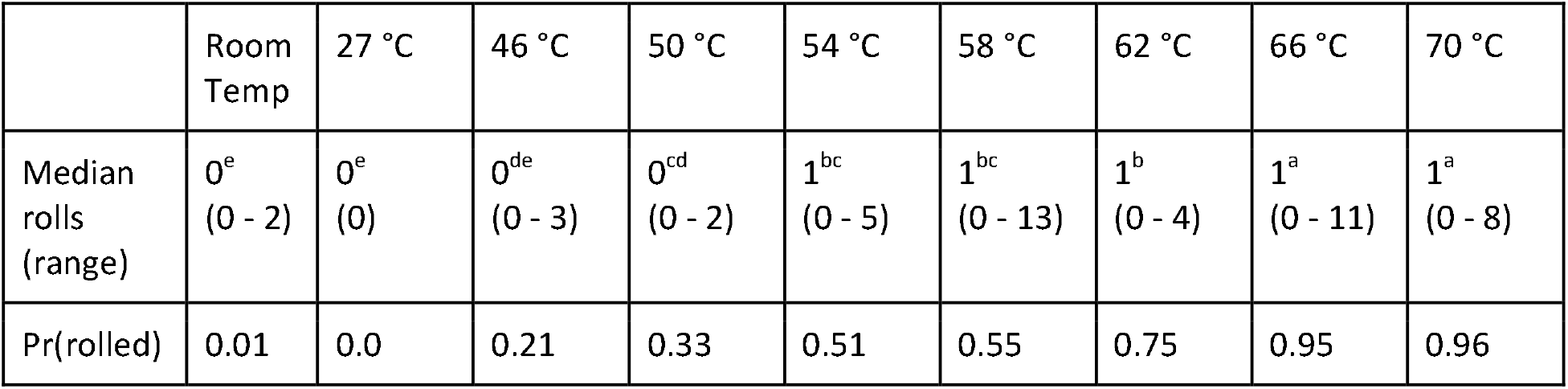
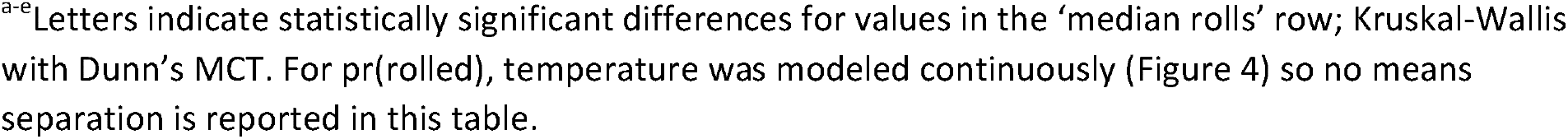
Median number of rolls and proportion of larvae that rolled at each tested temperature.

|  | Room Temp | 27 °C | 46 °C | 50 °C | 54 °C | 58 °C | 62 °C | 66 °C | 70 °C |
| --- | --- | --- | --- | --- | --- | --- | --- | --- | --- |
| Median rolls (range) | 0<sup>e</sup><br>(0 - 2) | 0<sup>e</sup><br>(0) | 0<sup>de</sup><br>(0 - 3) | 0<sup>cd</sup><br>(0 - 2) | 1<sup>bc</sup><br>(0 - 5) | 1<sup>bc</sup><br>(0 - 13) | 1<sup>b</sup><br>(0 - 4) | 1<sup>a</sup><br>(0 - 11) | 1<sup>a</sup><br>(0 - 8) |
| Pr(rolled) | 0.01 | 0.0 | 0.21 | 0.33 | 0.51 | 0.55 | 0.75 | 0.95 | 0.96 |

<sup>a-e</sup>Letters indicate statistically significant differences for values in the 'median rolls' row; Kruskal-Wallis with Dunn's MCT. For pr(rolled), temperature was modeled continuously (Figure 4) so no means separation is reported in this table.

**Figure 4.**
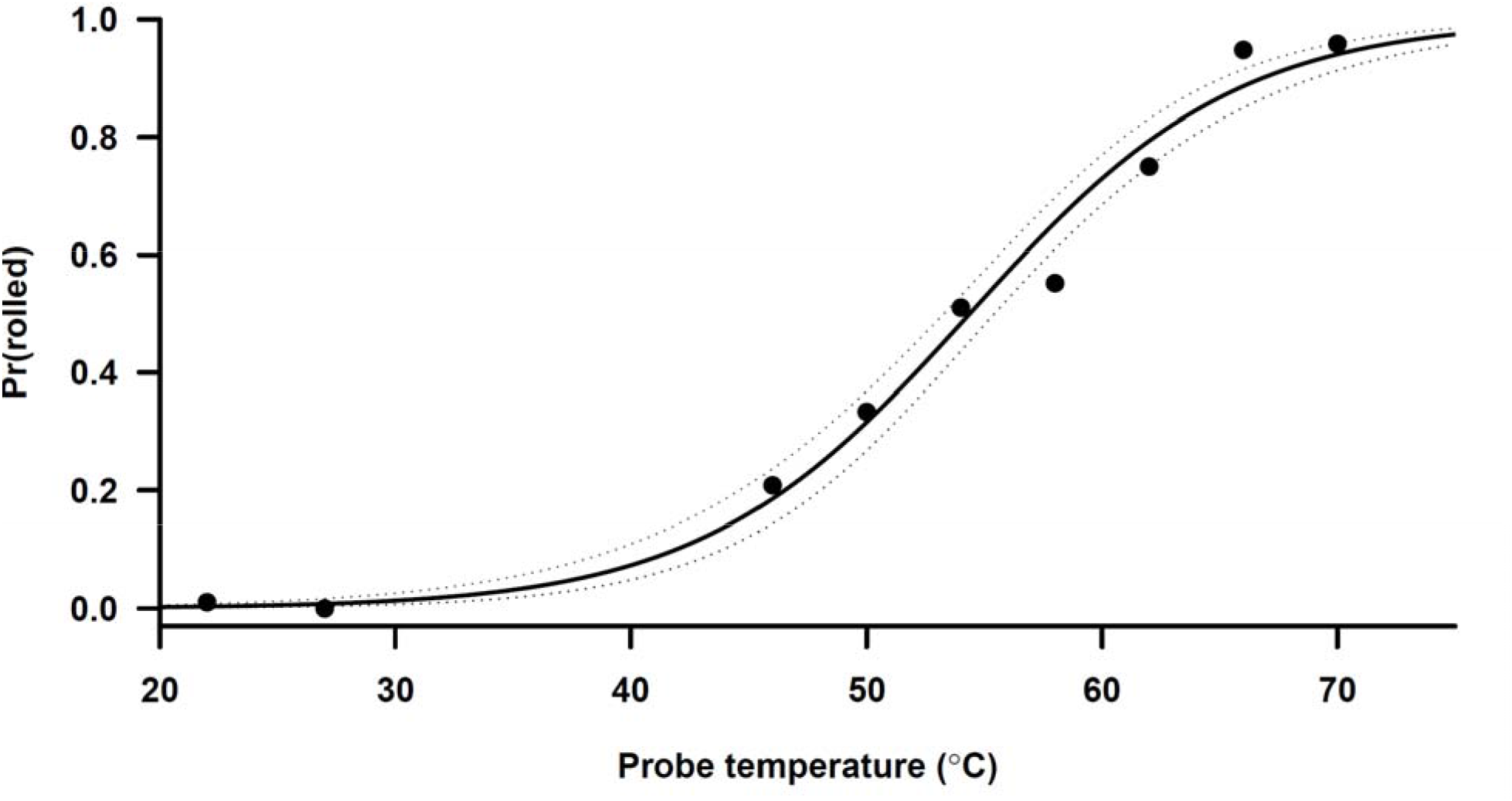
Proportion of L5/6 that rolled at each tested probe temperature at room temperature and set temperatures from 27 - 70 °C. Line of best fit shown 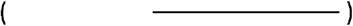 with 95% CI (dotted).

The number of rolls exhibited by a larva was also temperature-dependent (Figure 5; Kruskal-Wallis: K-W = 400.9, p < 0.0001), and also variable across larvae (0 - 13 rolls exhibited). At room temperature through 50 °C, the median was 0 rolls/larva; at 54 - 70 °C, the median was 1 roll/larva (Table 1).

**Figure 5.**
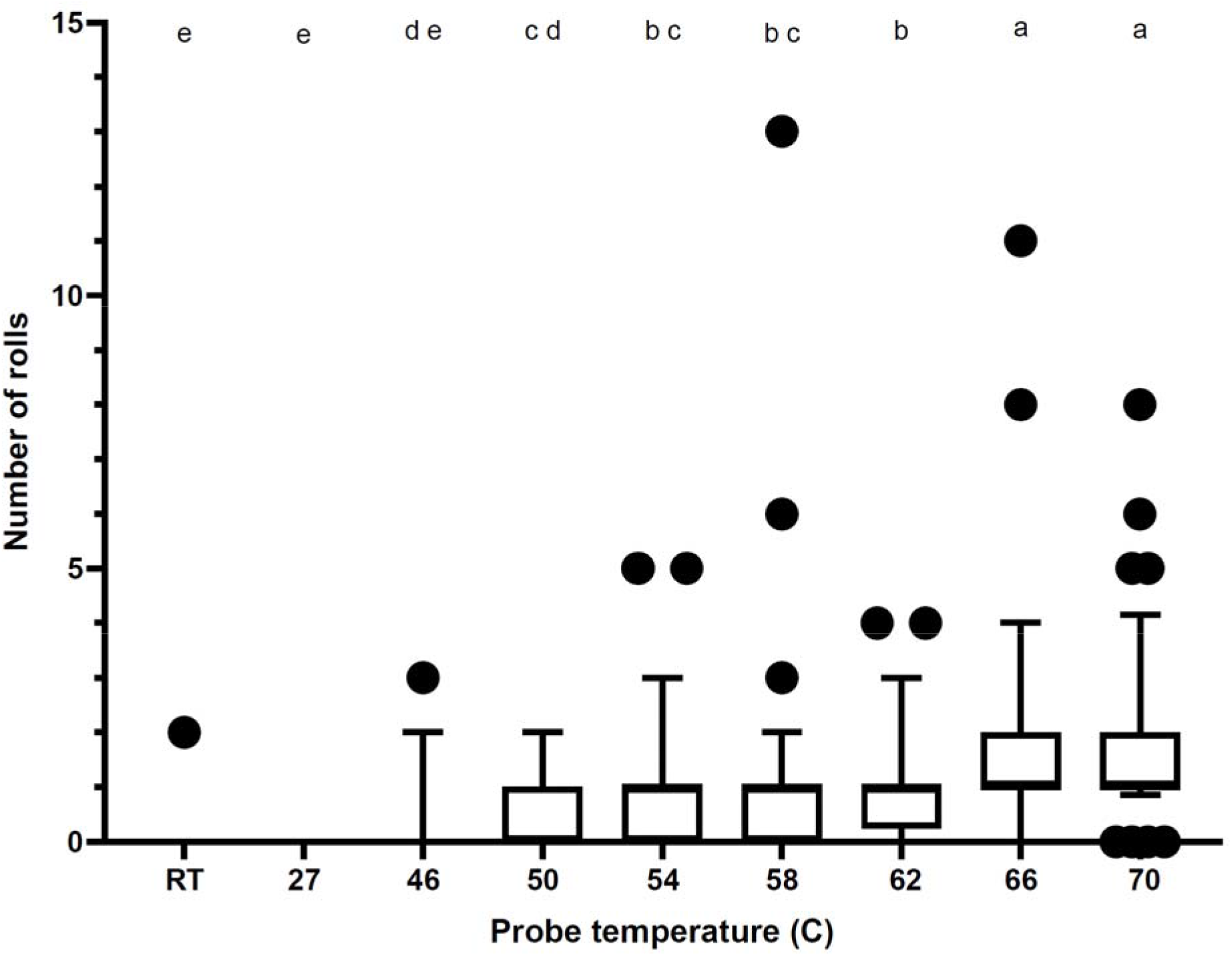
Medians (thick horizontal line) with 5-95% percentile (whiskers) of the number of larval rolls in response to probe temperature from room temperature (approximately 21 - 23 °C), and 27 - 70 °C set points. Letters indicate statistically significant differences among treatments (Dunn’s MCT; p < 0.05).

## Discussion

We found that BSFL exhibit nocifensive responses to thermally noxious stimuli, with localized responses occurring in over half the population by 54 °C and global responses occurring for the average L1 at 42.69 °C and average L5/6 at 39.70 °C. Late instar BSFL showed a more clear and consistent behavioral progression in the global thermal nociception assay, with obvious evidence of thrashing and generally more common contractions and longer holds compared to L1. The duration of exposure to the noxious stimulus while the larva was still able to control its behavior (the time between thrashing onset and seizure) correlated with the number of nocifensive curls observed. Importantly, rolling and general movement behaviors occur in the water and cannot be used to distinguish heated from unheated controls; however, thrashing, curling, contracting/holding, seizure, and paralysis are all responses unique to the heated treatments.

As with last instar *D. melanogaster* larvae, thrashing, seizing, and paralysis were most consistently displayed in late instar BSFL than the intermediate progression curling and contracting/holding behaviors. These behavioral data, combined with genetic data on the transient receptor potential (TRP) channels found in black soldier flies (Goldberg et al. 2024), make it likely that BSFL have functional ion channels responsive to high temperatures, as seen in closely-related dipterans and many other insects (Peng et al. 2015; Gibbons et al. 2022). The air bubble produced by L1 made it difficult for them to move and thus made assaying their nocifensive behaviors more challenging for the global assay (at least, until paralysis and seizure).

BSFL exhibited numerous differences from the nocifensive behaviors described for *D. melanogaster* in Chattopadhyay et al. (2012). For instance, BSFL did not tend to roll, and never whip, in the global thermal nociception assay. Instead, they showed a unique contract-and-hold response and frequent curling, with rolls occurring only incidentally during other behaviors as their bodies twisted. These differences could result from their different morphology - BSFL are broader and flatter than the more cylindrical *D. melanogaster* larvae (however, it is not clear why this would impact their ability to perform rolling behavior in water and BSFL *do* exhibit 360 degree lateral rolls when dry during the local nociception assay).

Another major difference is the temperature at which the nocifensive behaviors commence: BSFL had much higher thresholds than vinegar fly larvae in both assays. For instance, in the global heat assay, average late instar BSFL thrashed at 39.70 °C (compared to 26.6 °C in vinegar flies), seized at 48.50 °C (34.4 °C), and were paralyzed at 54.82 °C (37.1 °C), meaning that even the paralysis temperature was lower for vinegar fly larvae than the thrashing temperature in BSFL. These data are consistent with observations of BSFL tolerating very high temperatures. For instance, BSFL are known to thermoregulate in a density-dependent manner (Klammsteiner et al. 2025) towards a temperature just short of their thermal maxima that may be greater than 10 - 20 °C above ambient (Li et al. 2023). Li et al. (2023) recorded in-substrate temperatures of 46 °C for actively-feeding, high-density populations of BSFL, a temperature just below the average seizure threshold of 48.50 °C we observed. Larvae are known to exhibit escape behaviors in response to lethal larval overheating events (Barrett et al. 2023) and engage in constant motion, often described as fountaining, which is known to increase collective eating rate (Shishkov et al. 2019) but could also be explored as a heat management strategy for larvae at high densities. Therefore, the higher global noxious heat threshold for BSFL compared to *D. melanogaster* is biologically relevant (and see: Chia et al. 2018). However, the molecular underpinnings of this higher heat tolerance is not yet known and deserves further study.

Additionally, 100% of vinegar fly larvae responded within 20 s to the heat probe from 48-52 °C (Chattopadhyay et al. 2012) yet only ∼21-33% of BSFL responded in the same temperature range when tested with 15 s of continuous exposure. BSFL did not approach 100% responsiveness until 66-70 °C (the highest temperature on the probe) was applied for 15 seconds. BSFL exhibited increased likelihood of responsiveness with higher probe temperatures whereas *D. melanogaster* consistently stopped responding, even with 20 seconds of continuous stimulation, at temperatures above 52 °C. Further, *D. melanogaster* larvae exhibited more rolls at lower but still noxious temperatures whereas BSFL exhibited increased rolls at higher temperatures. As previously discussed, however, this could be a result of a difference in assay design, as BSFL were exposed to the stimulus for a constant amount of time while vinegar fly larvae were only exposed until they rolled away. Future research might attempt to quantify probe response latency in BSFL as ‘time to first attempted roll’ for larvae that are held under the probe, though it could be difficult to distinguish ‘rolling attempt’ from ‘general head movements’ that doesn’t precede a roll. If this were possible, the assay would prove more directly comparable to the vinegar fly literature. Alternately, differences in body size or evolutionary history between these species could explain these differences in nocifensive behavior.

Data from the global heat assay proves to be particularly relevant in the applied context for BSFL. Chattopadhyay et al. (2012) note that the global heating assay “…is more akin to the animal sitting in a heating cauldron… Although it is not clear when a larva might experience a globally noxious stimulus in the wild”. However, BSFL may experience this situation in two ways on farms: 1) when reared at such high densities that lethal larval overheating occurs (described in Barrett et al. 2023; Li et al. 2023), resulting in a globally noxious environment of high-temperature larvae/substrate on all sides; and 2) when boiled, blanched, scalded, or baked as a method of slaughter (Barrett et al. 2023; Sánchez-Velázquez and Hernández-Álvarez 2026).

These data suggest that larvae may respond nocifensively to temperatures above 39.70 °C and that death, measured by the proxy of paralysis, does not occur until the larvae reach a temperature of nearly 55 °C. Validating and adopting thermal slaughter standard operating procedures that minimize the duration of time larvae spend within the nocifensive window (above 39.70 °C but below the average paralysis temperature of ∼55 °C) would be essential to promoting larval welfare during slaughter (and see: Almeida et al. 2026). Producers can also use the nocifensive behaviors described in this protocol alongside already described escape behaviors to assess the potential for negative welfare states caused by exposure to thermally noxious stimuli during rearing and slaughter on their farms.

## Acknowledgments

We thank Edward Waddell and Nathan Fried for extensive conversations about black soldier fly larval nociception. We thank I Therese Veloso for helping to code videos of the global thermal nociception assay and Terry McGlynn for lending lab space to the same assay.

## Supplementary Figures

**Supplementary Figure 1.**
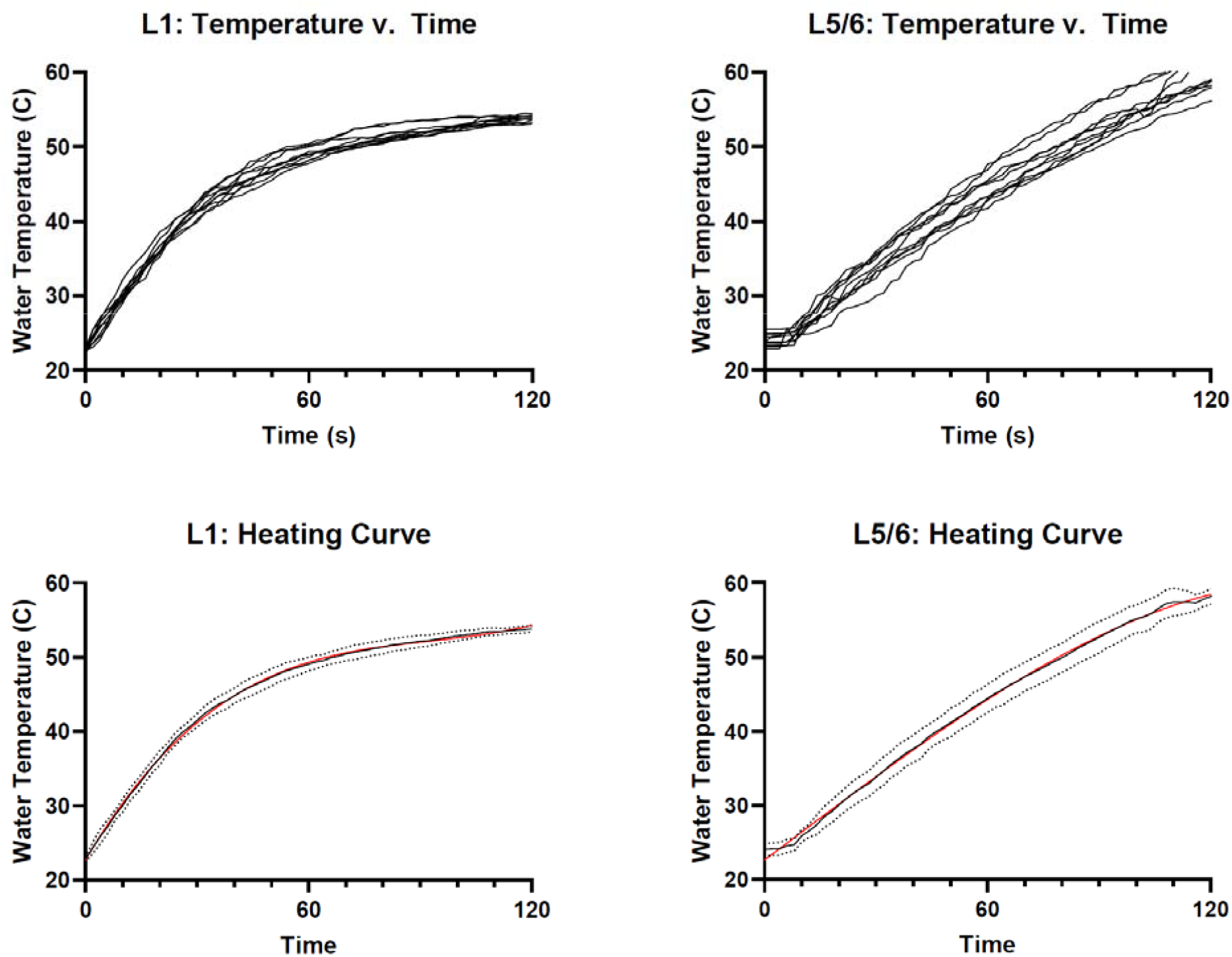
Temperature of water droplet vs. time for L1 and L5/6 BSF. Temperature of the water droplet in ten heating trials, when the water droplet is A) 50 μL on PTFE plate, hot plate set point at 60 °C (L1) and B) 1400 μL in 4 dram glass vial, hot plate set point at 115 °C (L5/6). C-D) Black lines show the average (dotted = 95% confidence interval) of the corresponding ten trials in A-B, with the red line demonstrating the best fit model for L1 (C: temperature = 3.05 e-05 (time^3^) -0.0085 (time^2^) + 0.84 (time) + 22.73; F = 925.5, df = 1, 606, R^2^ = 0.99, p < 0.0001) and L5/6 (D: temperature = -6.49 e-06 (time^3^) + 0.0001 (time^2^) + 0.38 (time) + 22.64; F = 13.38, df = 1, 607, R^2^ =0.98, p = 0.0003).

